# Profiling proteome-level amino acid substitutions in Alzheimer’s disease brain tissue

**DOI:** 10.64898/2026.09.08.750259

**Authors:** Peize Zhao, Shengzhi Lai, Shuaijian Dai, Chen Zhou, Ning Li, Weichuan Yu

**Affiliations:** Individualized Interdisciplinary Program, The Hong Kong University of Science and Technology, Hong Kong, China; Department of Electronic and Computer Engineering, The Hong Kong University of Science and Technology, Hong Kong, China; Division of Life Science, The Hong Kong University of Science and Technology, Hong Kong, China; Department of Biomedical Engineering, School of Basic Medical Sciences, Central South University, Changsha, Hunan Province, China

**Keywords:** amino acid substitution, TMT proteomics, Alzheimer’s disease, peptidoform identification, amino acid substitutomics

## Abstract

As the most prevalent neurodegenerative disorder worldwide, Alzheimer’s disease (AD) remains incompletely understood at the proteome level. In particular, current studies on amino acid (AA) substitutions have predominantly relied on genomic and transcriptomic profiling. Proteome-scale AA substitutions remain largely uncharacterized. Nevertheless, alterations at the DNA and RNA levels cannot fully recapitulate the spectrum of AA substitutions observed at the proteome level. This critical research gap persists largely due to the inherent analytical challenges posed by large-scale proteomic datasets. In this study, we address this limitation by analyzing two independent AD proteomic datasets, AMP-AD and PXD013753, using PIPI-C, an open-search mass spectrometry engine capable of resolving multiple co-occurring modifications per peptide.

We introduce a pipeline that enables the characterization of AA substitutions at the proteome level and the dissection of regulatory functions of key proteins with such substitutions. In both datasets, we observe that, after controlling for ambiguous post-translational modification mass shifts, the N>M substitution is the most frequent variant among N>X substitutions in the AD data. Furthermore, among the 701 overlapping proteins shared by the two datasets, we identified 6 literature-reported substitution sites, including residues 242 and 352 in glial fibrillary acidic protein, site 370 in actin gamma 1, as well as sites 111, 115, and 116 in hemoglobin subunit beta. Since most reported AA substitutions rely on genomic or transcriptomic evidence, and our pipeline adopts rigorous false positive control, those unreported substitutions are presumably detectable only via proteomic methods, emphasizing the unique value of our proteome-based substitutomics workflow for AD research.

## 1 INTRODUCTION

Alzheimer’s disease (AD) is a major neurodegenerative disorder affecting millions of individuals worldwide. The core clinical features of AD include progressive memory loss, executive dysfunction, and ultimately global cognitive decline [1]. These cognitive deficits are accompanied by prominent histopathological alterations, such as hippocampal atrophy and subsequent neuronal loss [2]. With the ongoing aging of the global population, the global economic cost associated with AD is anticipated to surge to USD 2 trillion by 2030 [1]. Despite extensive research efforts, the etiology of AD remains elusive, and existing treatments only offer symptomatic relief without halting disease progression. This highlights the urgent need to elucidate the molecular mechanisms underlying AD pathogenesis, which is important for the development of more effective therapeutic interventions.

A critical, yet incompletely defined driver of AD progression is the accumulation of amino acid (AA) substitutions in cellular proteins. Such modifications disrupt native protein structure and function. In turn, such alterations can compromise proteostasis and reinforce the pathological hallmarks of the disease [3, 4, 5]. AA substitutions originate from three distinct sources: DNA mutations, mRNA mutations, and errors during protein translation [6]. However, previous research has extensively focused on genomic [7, 8, 9] and transcriptomic alterations [10, 11, 12] to AD pathogenesis. In contrast, the majority of AD proteomic studies have focused heavily on post-translational modifications (PTMs) [13, 14, 15, 16]. Only a small fraction of research has targeted AA substitutions. Existing studies on AA substitutions in AD are confined to a small panel of pre-defined variants, leaving the global landscape of proteome-level substitutions largely unmapped. For example, the C410Y and L435F mutations in presenilin-1 (PSEN-1), as well as the N141I, T122P, M239V, and M239I mutations in PSEN-2, have been shown to alter the ratio of β-amyloid protein (Aβ) [17, 18]. However, this fragmented research approach constitutes a major knowledge gap: comprehensive, large-scale analyses of AA substitutions are essential for fully elucidating their roles in AD pathogenesis.

Nowadays, the comprehensive identification of AA substitutions in peptides poses substantial technical challenges in AD proteomic research, due to the inherent complexity of coexisting substitutions and the difficulty of balancing detection coverage with reliability. To address this key technical bottleneck, we use the AA substitutomic search pipeline that integrates open and closed search strategies to achieve both high coverage and robust validation of AA substitution events [19]. Specifically, the AA substitutomic pipeline employs PIPI-C [20] as the primary open-search engine, which is powerful for identifying non-specific AA substitutions, even in cases where multiple substitutions occur on the same peptide. Therefore, PIPI-C enables unbiased, global profiling of substitution events across the proteome. To mitigate false discoveries and ensure identification accuracy, all candidate substitutions are re-scored using Comet [21], a well-established closed-search engine widely recognized for its reliability in proteomic analysis, which requires a pre-specified list of variable modifications during the database search. Using this framework on the AMP-AD dataset, we systematically profile AA substitution patterns in two functionally important protein groups: cytosolic proteins and protein-binding proteins. Our analysis reveals distinct AA substitution signatures that correspond to specific protein functional perturbations, with clear relevance to neurodegenerative pathophysiology. We verify the biological significance of key substitutions through literature curation. To enhance the generalizability of our results, we analyze another independent AD dataset (PXD013753). The high replication rate observed across the two datasets supports the generalization of our findings.

## 2 RESULTS

### 2.1 AA substitutomic workflow and AA substitutions landscape of AD proteome

Previous studies have identified disease-relevant AA substitutions in AD-associated proteins [17, 18]. However, such investigations have been largely restricted to a small set of predefined AA substitutions and predominantly rely on genomic interrogation for variant detection. Here, we aimed to globally profile AA substitutions in AD from a proteomic standpoint, employing an unrestricted search strategy to capture all potential substitution events. We analyzed proteomic datasets from the AMP-AD consortium [22], encompassing superior temporal gyrus (STG) brain tissue specimens from 244 individuals across control and pathologically defined AD cases [23] (Figure 1A). All specimens were uniformly homogenized and analyzed via tandem mass tag-based mass spectrometry (TMT-MS).

**Figure 1:**
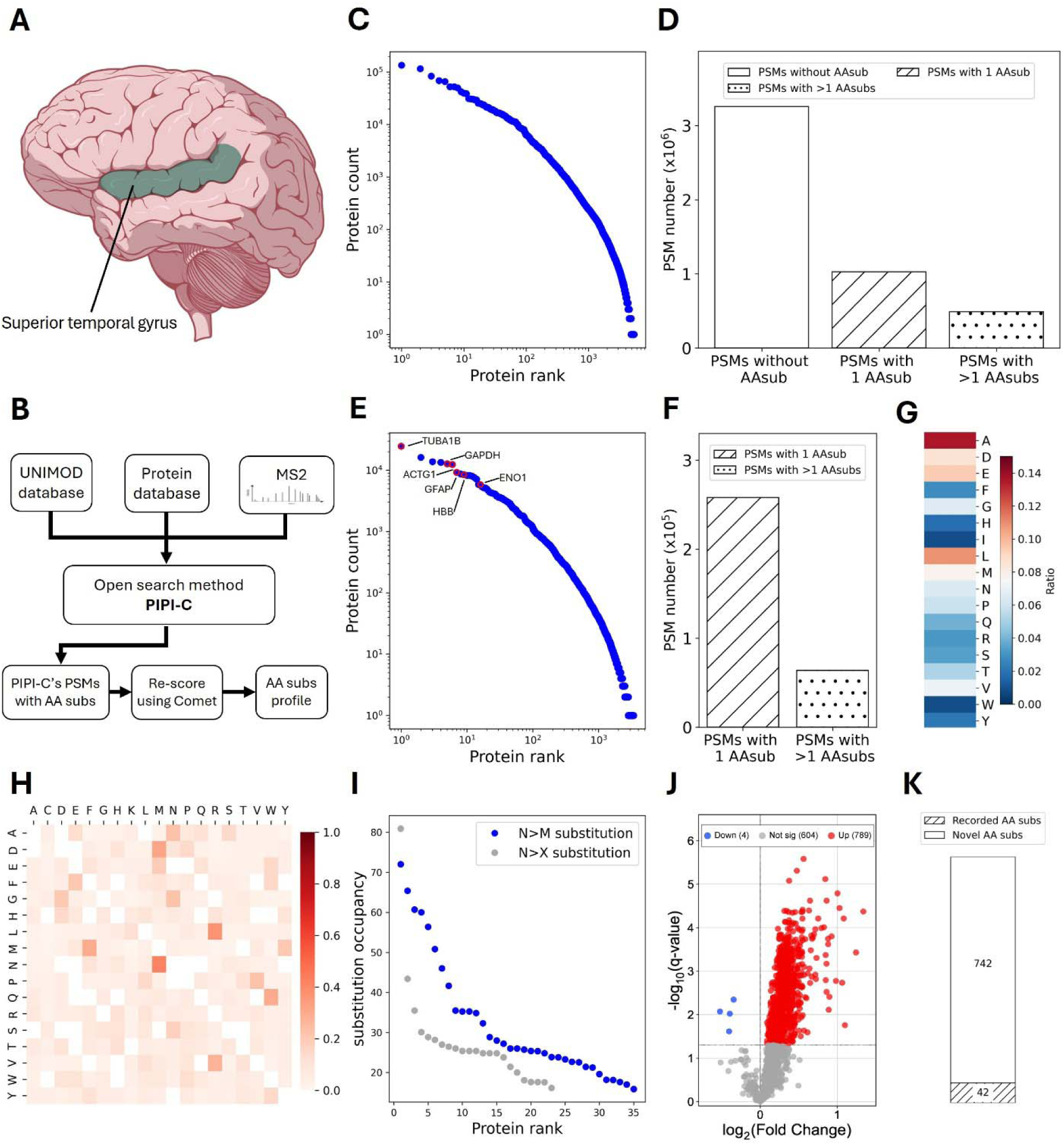
AA substitutomic analysis workflow and AA substitution landscape of AD. (A) The schematic representation of the human brain illustrates the location of the STG (green). (B) The workflow of proteomic dataset analysis. (C) Logarithm-logarithm scale distribution of the protein identification frequency of the PIPI-C-identified PSMs. (D) The identification results from PIPI-C. The clean section represents PSMs that lack any AA substitutions, the slashed section indicates PSMs containing a single AA substitution, and the dotted section denotes PSMs that have multiple AA substitutions. (E) Logarithm-logarithm scale distribution of the protein identification frequency of the Comet-validated identification results; the six proteins characterized in Sections 2.2.2 and 2.2.3 are marked in red. (F) The validated identification results from Comet. The slashed section indicates PSMs containing a single AA substitution, and the dotted section denotes PSMs that have multiple AA substitutions. (G) Normalized ratio of substituted AAs. (H) Heatmap displays the normalized ratio for all categories of AA substitutions in validated PSMs within the AMP-AD dataset. The vertical axis represents the original AAs and the horizontal axis denotes the AAs following substitution; each row is normalized to sum to one. I and L are reported together in this matrix. (I) Distribution of the N>M and N>X substitution occupancy in the AMP-AD dataset. Blue dots show the N>M substitution occupancy. Gray dots show the N>X substitution occupancy. (J) The volcano plot illustrates the quantification results for the quantified UPSPs that carry an unambiguous AA substitution. Up-regulated UPSPs are marked in red, down-regulated UPSPs in blue, and UPSPs without significant regulation in gray. The x-axis shows the log2 ratio of each UPSP relative to the common reference channel, and the dashed horizontal line marks q = 0.05. (K) The bar plot illustrates that 42 out of the 784 distinct AA substitutions have been documented in the UniProt protein variant database. Hatched: recorded AA subs; open: novel AA subs.

The AA substitutomic analytical workflow is outlined in Figure 1B. To achieve comprehensive profiling of diverse AA substitutions, open-search algorithms capable of simultaneous multi-modification detection are indispensable. We selected PIPI-C [20] as our core open-search engine, which detects multiple co-occurring modifications on individual peptides [19], a requirement for unbiased global profiling of AA substitutions. MS data derived from the AMP-AD proteomic datasets [22], together with the widely recognized modification and substitution library of UNIMOD [24], were utilized as the input for PIPI-C. To re-evaluate PIPI-C identifications under an orthogonal scoring framework, we re-searched the substitution-bearing spectra with Comet [21], a well-established closed-search engine. PIPI-C-derived modification profiles were supplied as variable modifications to Comet. Both engines were run with a PSM-level FDR cutoff of 1% controlled via target-decoy. Besides, we observed that mass shifts of certain AA substitutions overlap with those of other PTMs; thus, ambiguous substitution events were excluded from downstream analyses to eliminate confounding signals and ensure result fidelity (Table 1).

**Table 1:**
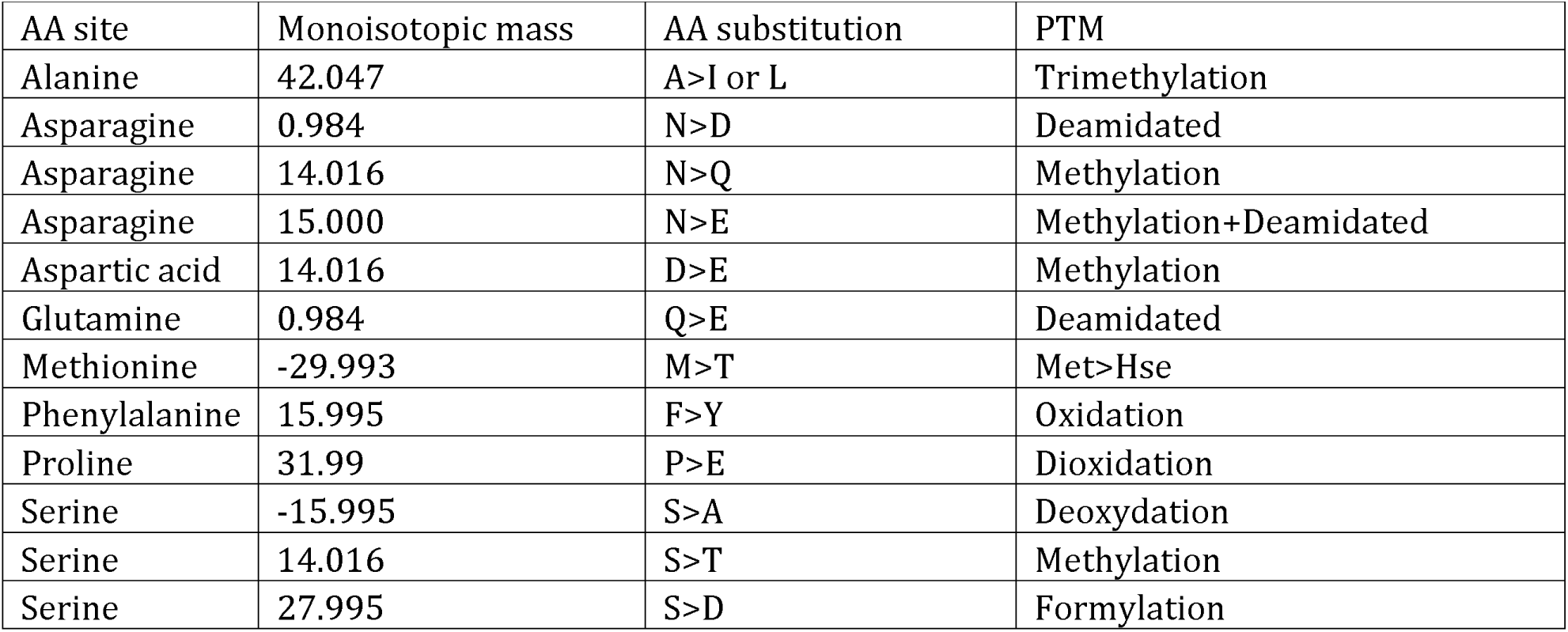
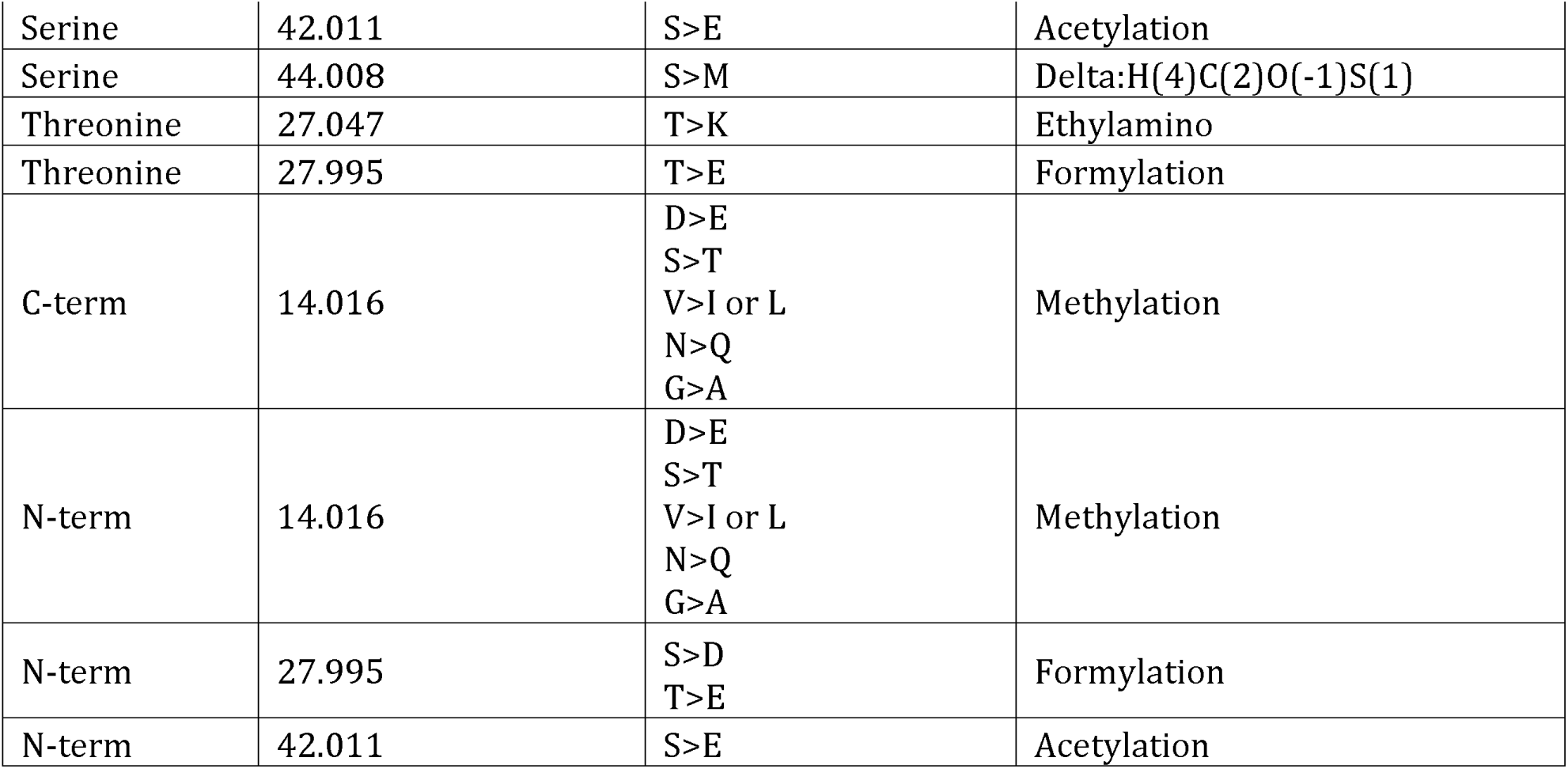
The ambiguous AA substitutions sharing monoisotopic mass shifts with common PTMs.

| AA site | Monoisotopic mass | AA substitution | PTM |
| --- | --- | --- | --- |
| Alanine | 42.047 | A>I or L | Trimethylation |
| Asparagine | 0.984 | N>D | Deamidated |
| Asparagine | 14.016 | N>Q | Methylation |
| Asparagine | 15.000 | N>E | Methylation+Deamidated |
| Aspartic acid | 14.016 | D>E | Methylation |
| Glutamine | 0.984 | Q>E | Deamidated |
| Methionine | -29.993 | M>T | Met>Hse |
| Phenylalanine | 15.995 | F>Y | Oxidation |
| Proline | 31.99 | P>E | Dioxidation |
| Serine | -15.995 | S>A | Deoxydation |
| Serine | 14.016 | S>T | Methylation |
| Serine | 27.995 | S>D | Formylation |
| Serine | 42.011 | S>E | Acetylation |
| Serine | 44.008 | S>M | Delta:H(4)C(2)O(-1)S(1) |
| Threonine | 27.047 | T>K | Ethylamino |
| Threonine | 27.995 | T>E | Formylation |
| C-term | 14.016 | D>E<br>S>T<br>V>I or L<br>N>Q<br>G>A | Methylation |
| N-term | 14.016 | D>E<br>S>T<br>V>I or L<br>N>Q<br>G>A | Methylation |
| N-term | 27.995 | S>D<br>T>E | Formylation |
| N-term | 42.011 | S>E | Acetylation |

The substitutomic pipeline generated robust identification results from the AD proteomic datasets. Figure 1C depicts the protein identification frequency distribution from PIPI-C: most proteins were detected 10 to 1,000 times. The relationship between protein identification frequency and rank follows Zipf’s law [25], consistent with the typical distribution of proteomic identifications. Figure 1D quantifies peptide spectrum matches (PSMs) from PIPI-C, categorized into non-substitution, single-substitution, and multi-substitution groups. After excluding PSMs attributed exclusively to TMT labeling mass shifts, about 31% of PIPI-C-identified PSMs carried a single AA substitution, and 15% harbored multiple substitutions. Retaining multi-substitution peptidoforms is the reason an open search was required here. PIPI-C identified substitution-associated spectra that were then extracted for validation by Comet. All validated PSMs contained at least one AA substitution. Figure 1E shows the protein frequency distribution of validated PSMs from Comet, which also conforms to Zipf’s law. Notably, about 20% of validated PSMs contained multiple AA substitutions (Figure 1F), demonstrating the widespread prevalence of substitution events in the AD proteome. Cysteine carbamidomethylation and lysine TMT labeling were set as fixed modifications, precluding substitution detection on these two residues; thus, substitutions were assessed across 18 AAs.

Substitution frequencies across these 18 residues were non-uniform. Expressed as a fraction of all unambiguous substitution events detected across the dataset, alanine was the most frequently substituted residue (about 14%), while isoleucine and tryptophan were the least frequent (each about 0.8%; Figure 1G). A heatmap of normalized substitution type ratios in the dataset (Figure 1H) revealed that the N>M substitution was the predominant variant among all N>X substitutions, accounting for about 42% of events (Supplementary File 1). To quantify the relative abundance of N>M substitutions, we calculated the N>M and N>X substitution occupancy for proteins identified in more than 60% of experiments, using the biotin occupancy-like metric [26, 27] defined in the Methods (Figure 1I, Supplementary File 2). N>M substitution occupancy exceeded nearly all other N>X variants, indicating that asparagine is preferentially converted to methionine relative to other AAs. All raw PIPI-C identification data and validated results are publicly archived in Zenodo [28] with record number 15010408.

Because open-search identifications of AA substitutions can be confounded by co-eluting modifications and by isotope-envelope errors, we manually inspected the fragment-ion evidence for representative substitutions in the proteins characterized below. Eight representative sites are shown in Supplementary Figure S1: GAPDH M175F, HBB A116Q, ACTG1 V370R, ENO1 V152T, GFAP Y242T, HBB L111P, GFAP L352R, and HBB L115A, with precursor mass errors within 8 ppm. Six of these sites, GFAP 242, GFAP 352, ACTG1 370, HBB 111, HBB 115, and HBB 116, are the positions reported in the literature and referred to in the Abstract; the three HBB sites lie on the same tryptic peptide, so a single spectrum localizes neighbouring events. Identifications were controlled at 1% PSM-level FDR and required agreement between two search engines. Because protein-level error rates are not bounded by PSM-level FDR, we do not attach a protein-level false-discovery estimate to the 185 proteins; the manually annotated spectra in Supplementary Figure S1 provide direct fragment-ion evidence for the substitutions discussed below.

We next quantified the differential regulation of AA substitutions with unique PTM site patterns (UPSPs) in AD samples relative to the common reference (CR) channel that is included in every TMTpro experiment (Figure 1J, Supplementary File 3). This analysis tests whether the abundance of each UPSP deviates significantly from the common reference across experiments; it is not a direct case-control contrast. Detailed quantification protocols and filtering criteria are described in the Methods section. Quantification revealed a striking predominance of UPSPs elevated relative to the common reference (789) over those reduced relative to it (4). The set of 789 UPSPs should be read as substitution events that are reproducibly detected; which of them track AD neuropathology is established separately in Section 2.2. These 789 upregulated UPSPs map to 784 distinct protein-level AA substitution events distributed over 185 proteins; identical substitutions detected on different peptidoforms of the same protein were counted once (Supplementary File 4). Of these 784 substitution events, 42 (5.4%) overlapped with germline/somatic variants curated in UniProt [29]; the remaining 742 events are not represented in UniProt and therefore correspond to proteome-level observations that may reflect either post-translational events or as-yet-unannotated nucleotide variants (Figure 1K, Supplementary File 4). AA substitution profiles varied across proteins: hemoglobin subunit beta (HBB) exhibited the highest normalized substitution ratio, with 22 substitution sites detected in its 147-AA sequence. Actin gamma 1 (ACTG1) carried the highest absolute number of substitutions, with 45 distinct substitutions identified. In subsequent sections, we focus on significantly regulated UPSPs, with an emphasis on upregulated AA substitutions and proteins harboring multiple substitution sites.

### 2.2 Multiple significantly up-regulated AA substitutions in AD biomarkers

Using our AA substitutomic analysis pipeline, we identified a total of 185 proteins carrying AA substitutions from the AMP-AD dataset (Supplementary Files 3 and 4). To prioritize functionally meaningful candidates, we applied two selection criteria: (1) proteins harboring multiple reproducibly quantified AA substitution events; (2) proteins with well-established and strong relevance to AD pathogenesis. By filtering the full list of 185 proteins based on these two rules, we finally narrowed down to six high-confidence AD-associated biomarker proteins. In addition, Gene Ontology (GO) enrichment analysis [30] demonstrated that these six core proteins were enriched in cytosol components and protein-binding functions. Comprehensive characterizations of the AA substitutions in these proteins are provided in the subsequent sections.

#### 2.2.1 Case-control analysis of AA substitutions in AD versus neuropathological controls

To establish which of these proteins carry substitutions that track AD neuropathology directly, rather than only being reproducibly quantified against the internal reference, we performed a case-control analysis using the harmonized neuropathological classification distributed with the AMP-AD dataset. Because every experiment contains both AD and control channels, the contrast was computed within each experiment on identical spectra, separately within Emory and within Mayo Clinic specimens, and the two estimates were combined by inverse-variance fixed-effect meta-analysis (Methods). Of the 1,678 substitution-bearing peptidoforms testable in both institutions, 138 substitution events in 52 proteins are more abundant in AD than in neuropathological controls at q < 0.05 with a positive effect in both institutions, and 142 substitution events in 75 proteins are less abundant (Supplementary File 6).

The elevated events are markedly concentrated rather than distributed across substituted peptidoforms as a class: the five largest contributors account for half of them, and four of these five, GFAP, GAPDH, ENO1, and hemoglobin subunit beta (HBB), are among the six proteins characterized below. Taken as a group, the 209 substitution-bearing peptidoforms of the six proteins rank well above the remainder in the AD-minus-control ordering, and this ordering is unchanged when peptidoforms are stratified by reporter-ion intensity. Individually, 91% of the substitution-bearing peptidoforms of GFAP, 92% of those of ENO1, 90% of those of HBB, and 90% of those of GAPDH are more abundant in AD, whereas ACTG1 contributes a single event to the 138 and TUBA1B carried no peptidoform testable in both institutions; both were prioritized on substitution density and structural relevance rather than on differential abundance. These differences reflect the disease-associated abundance of the substituted proteoforms themselves; because reporter-ion ratio compression is intensity-dependent and substitution-bearing peptidoforms are systematically less intense than unmodified ones, the present data do not resolve whether substitution stoichiometry per molecule of protein also changes.

#### 2.2.2 AA substitutomes of cytosol-related proteins in the AD proteome

Cytosol, also termed the cytoplasmic matrix, is an essential liquid component of eukaryotic cells [31]. The core function of the cytosol is to mediate the transport of metabolites from their biosynthetic sites to their functional utilization sites [32]. Multiple studies have established associations between cytosolic proteins and AD pathogenesis. For instance, Aβ accumulates in the cytosol of lens fiber cells in AD patients [33]. Furthermore, aberrant hyperphosphorylation of the tau protein in the cytosol impairs tau-mediated microtubule stabilization, thereby driving neuronal degeneration and cognitive decline in AD [34].

In this study, GO enrichment analysis revealed significant enrichment of the cytosol term in the AD proteomic dataset. Among cellular component terms enriched for upregulated proteins, the cytosol was one of the most highly represented categories, encompassing 118 target proteins (Figure 2A). These enrichment results highlight a critical direction for investigating the link between AA substitutions and cytosol-related proteins in the AD proteome. Besides the cytosol, extracellular exosomes were also significantly enriched in upregulated proteins of the AD proteomic dataset.

**Figure 2:**
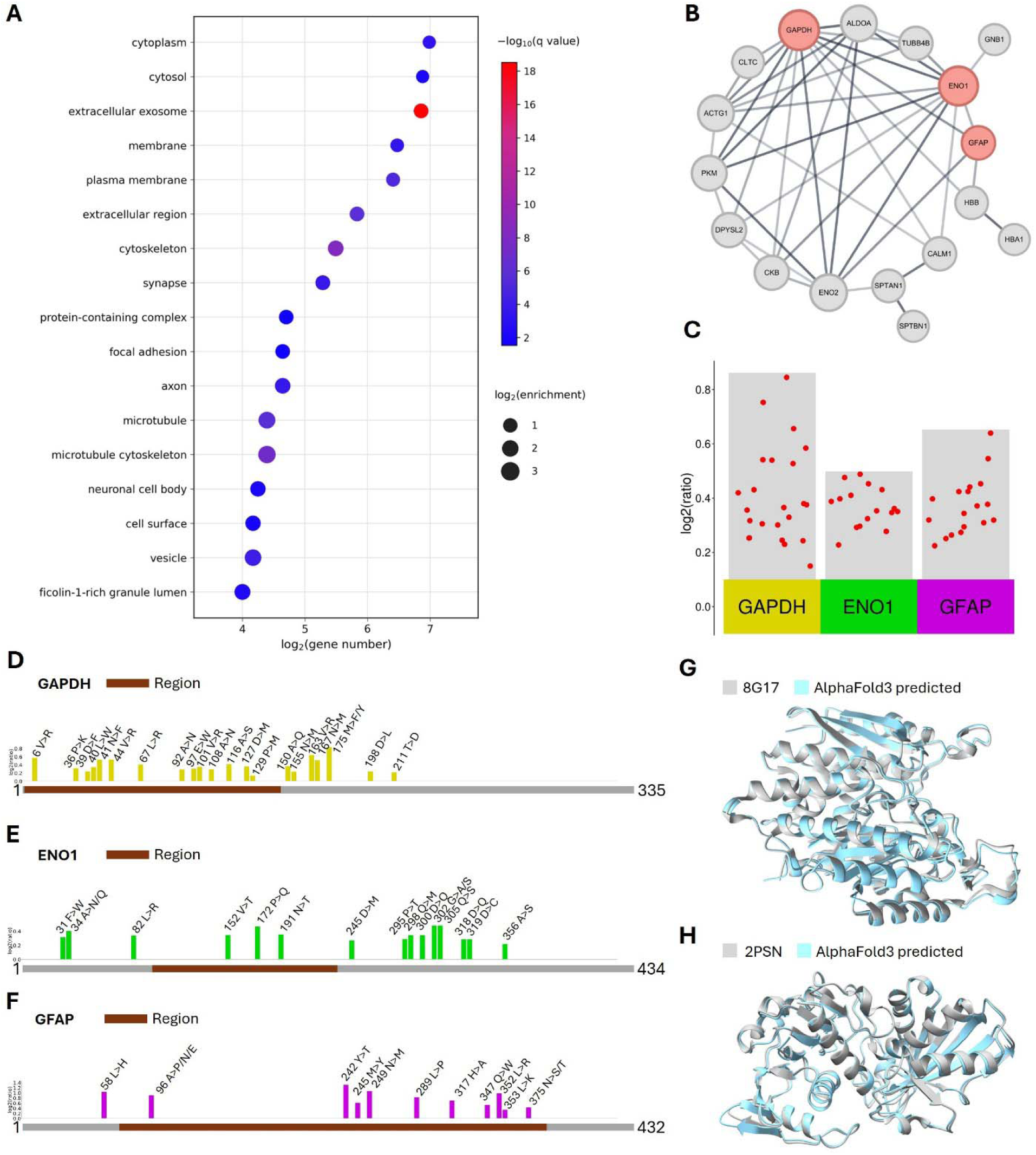
AA substitutomes of cytosol-related proteins in the AD proteome. (A) Bubble plot of the GO cellular component enrichment from the upregulated results. The size and color of the bubble represent the enrichment fold of the genes and their minus base 10 logarithm of the q-value, respectively. (B) Network of 17 proteins associated with the cytosol, where the size of each node reflects the degree of the respective protein. (C) Volcano plot displaying the quantification results for the up-regulated UPSPs of these proteins. (D) Cumulative AA substitutions map of GAPDH. Brown segment: binding and interaction region. (E) Cumulative AA substitutions map of ENO1. Brown segment: binding and interaction region. (F) Cumulative AA substitutions map of GFAP. Brown segment: structural domain. (G) Structural comparison is conducted between the cryo-electron microscopy structure of human GAPDH (PDB: 8G17) and the structure predicted based on cumulative AA substitutions. The structure of 8G17 is shown in gray, while the AlphaFold3-predicted structure incorporating cumulative AA substitutions is shown in light blue. (H) Structural comparison is conducted between the crystal structure of human ENO1 (PDB: 2PSN) and the structure predicted based on cumulative AA substitutions. The structure of 2PSN is shown in gray, while the AlphaFold3-predicted structure incorporating cumulative AA substitutions is shown in light blue.

Figure 2B displays the protein-protein interaction (PPI) network of 17 cytosolic proteins identified with at least 10 AA substitutions in the dataset. Node size in the network corresponds to the connectivity degree of each protein. Among these proteins, glyceraldehyde-3-phosphate dehydrogenase (GAPDH) exhibited the highest connectivity degree in the PPI network. GAPDH is a key glycolytic enzyme that catalyzes glucose breakdown to generate energy and carbon metabolites. In addition to its canonical role in aerobic glucose metabolism, emerging evidence has implicated GAPDH in diverse non-metabolic processes. Notably, neuroproteomic studies have demonstrated robust interactions between GAPDH and core AD-related proteins, including Aβ, amyloid precursor protein, and tau [35, 36].

In addition to GAPDH, two other pivotal proteins, alpha-enolase (ENO1) and glial fibrillary acidic protein (GFAP), are also extensively characterized as AD-associated proteins. Based on our quantitative results, we selected up-regulated UPSPs and visualized them in Figure 2C. The specific up-regulated AA substitutions identified in these three proteins are detailed in Figure 2D–F. For GAPDH, we mapped 21 AA substitution sites from the AD proteomic dataset (Figure 2D, Supplementary File 3). Notably, at site 175, methionine is substituted by either phenylalanine or tyrosine. Residues 2–148 of GAPDH interact with one isoform of tryptophanyl-tRNA synthetase (WARS1), a domain essential for regulating and mediating the aminoacylation process [37]. Within this region, we identified 14 upregulated substitution sites; whether these substitutions functionally affect WARS1 binding requires biophysical validation.

ENO1, a glycolytic enzyme isoform, catalyzes the conversion of 2-phosphoglycerate to phosphoenolpyruvate [38] and also carries multiple up-regulated AA substitutions. ENO1 is a well-documented target of oxidative modification in the brains of AD patients [39] and is detectable across all three pathological stages of AD and in multiple AD models [40, 41]. Our identification results are consistent with these previous reports: we mapped 15 AA substitution sites in ENO1 from the AD proteome (Figure 2E, Supplementary File 3). Notably, at site 34, alanine is substituted by asparagine or glutamine; at site 302, glycine is substituted by alanine or serine. ENO1 contains a critical functional region (residues 97–237) required for repressing MYC promoter activity [29]. Within this region, we identified three up-regulated AA substitutions: V152T, P172Q, and N191T. Previous studies report MYC up-regulation in degenerating neurons of traumatic injury animal models [42], and MYC dysregulation is pathologically linked to AD and other neurodegenerative disorders [43]. The three substitutions in the 97–237 region may attenuate the MYC repressive function of ENO1, a possibility that requires functional testing.

GFAP, an astrocytic cytoskeletal protein measurable in peripheral blood, was associated with numerous AA substitutions in our study. It plays an indispensable role in maintaining the mechanical strength and morphological integrity of astrocytes [44]. Recent studies validate GFAP as a reliable indicator of AD pathology, with strong potential for early disease diagnosis [45, 46]. We identified 11 up-regulated AA substitution sites in GFAP (Figure 2F, Supplementary File 3), and the elevated abundance of GFAP is consistent with its established role as an AD biomarker. GFAP further accounts for 17 of the 138 substitution events that are more abundant in AD than in neuropathological controls in both contributing institutions (Supplementary File 6), making it the protein with by far the largest mean effect size in that set and the largest contributor under the stricter within-institution criterion. Notably, 10 of these substitutions localize to the intermediate filament rod domain (residues 69–377) [29], a region critical for protein structural integrity. The high substitution frequency in this domain may disrupt the coiled-coil rod structure, impairing GFAP stability and function. At site 96, alanine is replaced by proline, asparagine, or glutamic acid; at site 375, asparagine is substituted by serine or threonine. Additionally, two substitution sites (242 and 352) are associated with Alexander disease, a rare neurodegenerative disorder. The Y242D substitution in GFAP impairs protein oligomerization and solubility by altering residue charge [47]; our identified Y242T substitution alters the residue side chain in a manner that may parallel the documented Y242D effect. The L352P substitution causes severe Alexander disease because proline acts as a helix breaker within the rod domain: its pyrrolidine ring restricts the backbone phi angle, and its imino nitrogen cannot donate the i to i+4 backbone hydrogen bond that stabilizes the alpha-helix [48]. In our study, we detected an L352R substitution, where hydrophobic leucine is replaced by hydrophilic arginine. This substitution may further reduce side-chain hydrophobicity at L352, indicating a more deleterious effect on GFAP function.

To evaluate the structural impacts of the identified AA substitutions, we compared the experimental structures of GAPDH [49] and ENO1 [50] with their AlphaFold3-predicted structures harboring cumulative AA substitutions (Figure 2G and H). This comparison reveals potential structural perturbations; the precise molecular consequences and functional impacts of these substitutions on GAPDH and ENO1 require further characterization.

#### 2.2.3 AA substitutomes of protein-binding proteins in the AD proteome

Protein binding is crucial in the pathological progression of AD. For instance, the binding affinity of tau to tubulin is regulated by its phosphorylation status, which is tightly modulated by the balanced activity of kinases and phosphatases [51]. Aberrant hyperphosphorylation of tau reduces its tubulin-binding capacity, triggering microtubule disassembly and neuronal dysfunction [52]. Additionally, characterization of the tau-fyn binding interface supports the development of small-molecule inhibitors targeting this interaction, representing a promising therapeutic strategy for AD [53]. Furthermore, studies have confirmed that binding of A*β* and tau oligomers to the amyloid precursor protein (APP) is required for APP internalization into neurons, leading to synaptic dysfunction and memory impairment [54]. Collectively, this evidence demonstrates that dysregulated protein binding is a key driver of AD pathogenesis.

In this study, GO enrichment analysis of the up-regulated UPSPs against the identified proteome reveals that the proteins are enriched in a range of specific protein-binding functions (Figure 3A). AA substitutions can alter protein primary structure, modifying protein folding and intra-molecular bonds, ultimately affecting binding sites and functionality [55]. The enrichment of specific binding terms, including identical protein binding, protein kinase binding, and protein domain-specific binding (Supplementary File 7), motivates the AA substitutomic investigation; the generic parent term protein binding is not itself enriched against the identified proteome. Among protein-binding-related proteins, the top three with the highest number of identified AA substitutions are ACTG1, HBB, and tubulin alpha-1B chain (TUBA1B) (Figure 3B). Additionally, based on our quantification results, the up-regulated UPSPs of the three proteins that have the most abundant AA substitutions are presented in Figure 3C.

**Figure 3:**
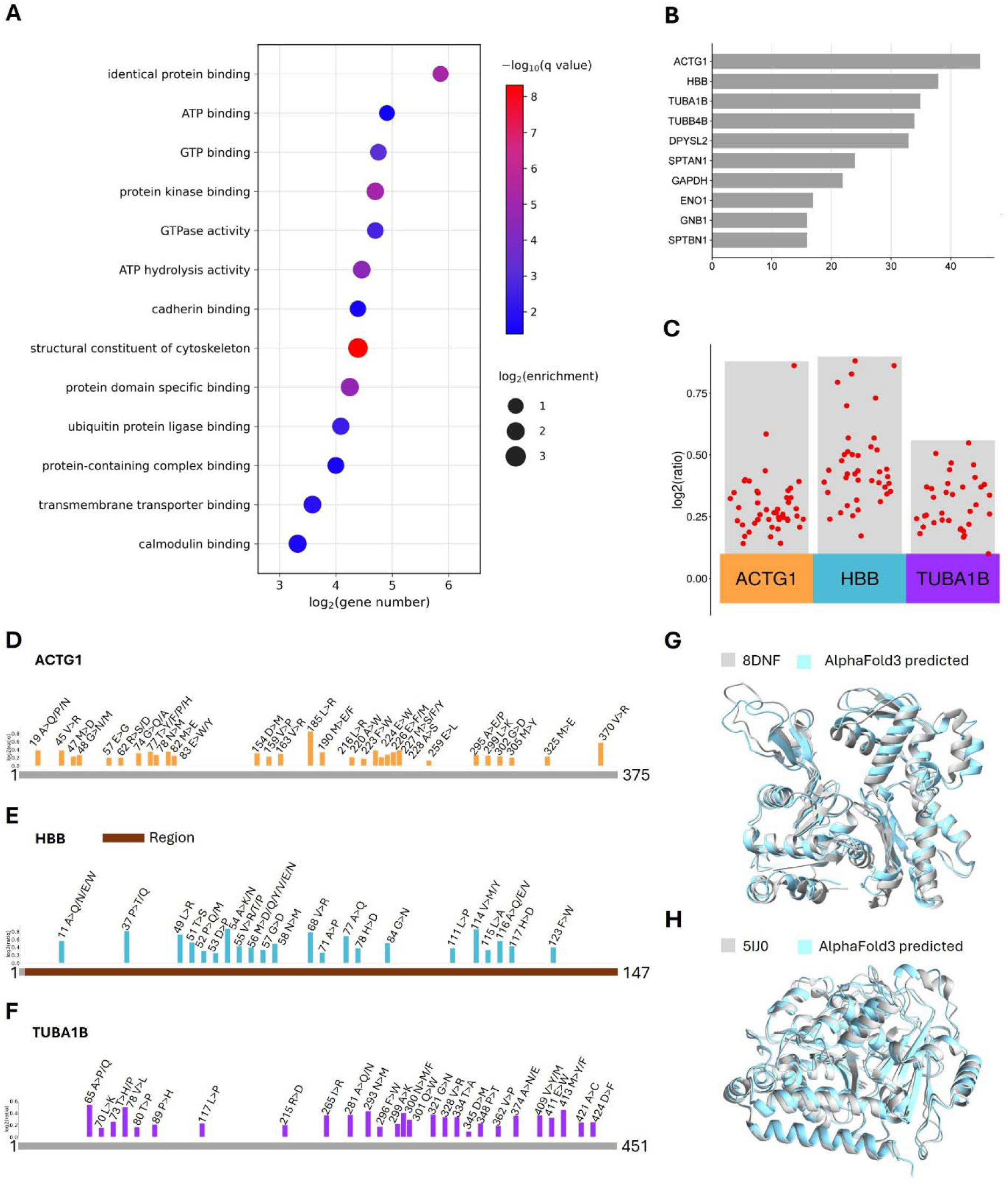
AA substitutomes of protein-binding proteins in the AD proteome. (A) Bubble plot of the GO molecular function enrichment from the up-regulated results. The size and color of the bubble represent the enrichment fold of the genes and their minus base 10 logarithm of the q-value, respectively. (B) Bar plot of the number of identified AA substitution events within the top ten proteins associated with protein binding. (C) Volcano plot presenting the quantification results for the up-regulated UPSPs of these proteins. (D) Cumulative AA substitutions map of ACTG1. (E) Cumulative AA substitutions map of HBB. Brown segment: structural domain. (F) Cumulative AA substitutions map of TUBA1B. (G) Structural comparison is conducted between the cryo-electron microscopy structure of human ACTG1 (PDB: 8DNF) and the structure predicted based on cumulative AA substitutions. The structure of 8DNF is shown in gray, while the AlphaFold3-predicted structure incorporating cumulative AA substitutions is shown in light blue. (H) Structural comparison is conducted between the cryo-electron microscopy structure of human TUBA1B (PDB: 5IJ0) and the structure predicted based on cumulative AA substitutions. The structure of 5IJ0 is shown in gray, while the AlphaFold3-predicted structure incorporating cumulative AA substitutions is shown in light blue.

ACTG1 is ubiquitously expressed in most cell types as a core cytoskeletal component that mediates intracellular motility. Emerging evidence has also linked ACTG1 variation to neurotoxicity; for example, carriers of the A allele of the synonymous ACTG1 variant rs1135989 (Ala310) show a markedly increased risk of high-grade vincristine-induced peripheral neurotoxicity [56]. A recent study also reported a positive correlation between ACTG1 expression levels and cognitive scores in AD patients [57]. In our analysis, ACTG1 exhibited the highest number of AA substitutions (45 substitution events distributed over 30 sites; Figure 3D, Supplementary File 4). Notably, eleven residue sites carried alternative AA substitutions, indicating that ACTG1 isoforms with distinct substitution combinations may contribute to AD phenotypic heterogeneity and warrant further investigation. Among these sites, residue 370 is a documented missense mutation locus in ACTG1. Functional assays showed that the V370A mutation impairs cell proliferation under thermal or hyperosmotic stress. Molecular modeling revealed that this mutation disrupts a protein-protein interaction interface in ACTG1, destabilizing the C-terminal tail and cytoskeletal networks [58]. The V370 residue maintains ACTG1 structural integrity via hydrophobic interactions with R116, V134, H371, and C374, interactions conserved across species to stabilize the ACTG1 C-terminus [59, 60, 61]. However, the V370R mutation introduces a hydrophilic residue that further disrupts C-terminal stability. Importantly, V370R shows one of the highest up-regulation ratios in ACTG1 UPSPs, suggesting it is among the most consistently detected ACTG1 substitutions in this cohort; its functional consequences warrant follow-up.

HBB was among the most extensively altered proteins in our study, with 22 distinct AA substitution sites distributed across its 147-residue protein chain, representing one of the highest substitution densities. All substitutions localized to the highly conserved globin domain (residues 3–147; Figure 3E, Supplementary File 3), which is essential for protein folding and oxygen-binding activity. The clustering of AA substitutions within this critical domain suggests profound effects on protein stability and molecular interactions. Eight residues (11, 37, 52, 54, 55, 56, 114, and 116) displayed multiple alternative substitutions. This high degree of sequence variability in AD patients suggests potential pathological dysregulation of HBB structure and function, offering new insights into its non-erythroid functions in neuronal homeostasis and AD pathogenesis [62]. We identified two dense substitution clusters: residues 51–58 and 114–117, with complete substitution coverage of all sites in these regions. These contiguous segments likely correspond to critical functional regions mediating subunit interactions, heme binding, or allosteric regulation, representing hotspots of AD-related molecular dysfunction.

In HBB, from the 22 identified AA substitution sites, three have been previously reported in the literature, specifically at positions 111, 115, and 116. The rest of the identified AA substitution sites remain unreported in existing studies. Since the majority of documented substitutions are discovered through genomic and transcriptomic profiling, these novel events can presumably be exclusively captured by proteomic analysis. Residues 115 and 116 lie within the 114–117 substitution cluster adjacent to site 111. Nucleotide mutations encoding these AA substitutions cause severe thalassemia phenotypes via hemoglobin dysfunction [63, 64, 65, 66]. Although pathogenic HBB variants are relatively rare [67], all three substitutions are linked to hemoglobinopathies characterized by moderate-to-severe microcytic anemia, indicating potential synergistic effects. These mutations destabilize the heme moiety by altering side chains within the heme pocket and localize to the G-helix core, severely impairing helical integrity [68]. Mounting evidence connects thalassemia to AD, including learning and memory deficits in thalassemic mouse models [69]. A nationwide retrospective cohort study further reported a 1.88-fold increased risk of dementia in thalassemia patients after adjusting for covariates [70]. Our identification of these AA substitutions at critical structural positions provides a molecular framework for elucidating the mechanistic link between hemoglobinopathies and AD pathogenesis.

TUBA1B, an essential *α*-tubulin isotype governing microtubule dynamics and cytoskeletal integrity, harbored 28 distinct AA substitution sites in our analysis (Figure 3F, Supplementary File 3). Notably, residues 65, 73, 281, 300, 374, 409, and 413 carried multiple alternative UPSPs, placing TUBA1B among the proteins with the highest polymorphic substitution density. As a core component of neuronal microtubules, TUBA1B maintains microtubule stability, axonal transport, and neuronal communication, functions severely compromised in AD [71]. Microtubule networks are extensively destabilized in AD, partially due to impaired structural integrity of TUBA1B-containing microtubules, directly driving synaptic degeneration and neuronal apoptosis [72, 73]. TUBA1B dysfunction is also tightly linked to tau pathology: under physiological conditions, tau stabilizes microtubules via direct binding to TUBA1B, while pathological tau in AD loses this binding affinity, causing defective mitochondrial trafficking, synaptic vesicle dysregulation, and accelerated neurodegeneration [72]. All TUBA1B UPSPs detected here are upregulated relative to the common reference. Independently of our quantification, recent studies report that elevated TUBA1B expression in AD-associated immune cells promotes pro-inflammatory cytokine secretion, exacerbating neuroinflammation and microtubule destabilization [74].

To evaluate the structural impacts of AA substitutions on protein-binding proteins, we compared the experimental structures of ACTG1 and TUBA1B with their AlphaFold3-predicted structures harboring cumulative substitutions (Figure 3G and H). While this analysis reveals potential structural perturbations, the detailed molecular consequences and functional impacts of these substitutions require further investigation.

### 2.3 Verification using one additional AD proteomic dataset

In preceding sections, various AA substitutions have been identified within the AMP-AD dataset. To assess the generalizability of our pipeline and qualitative replication of substitution-site recurrence, we use another AD dataset. This additional dataset is obtained from ProteomeXchange [75] under the data identifier PXD013753 [76].

**Figure 4:**
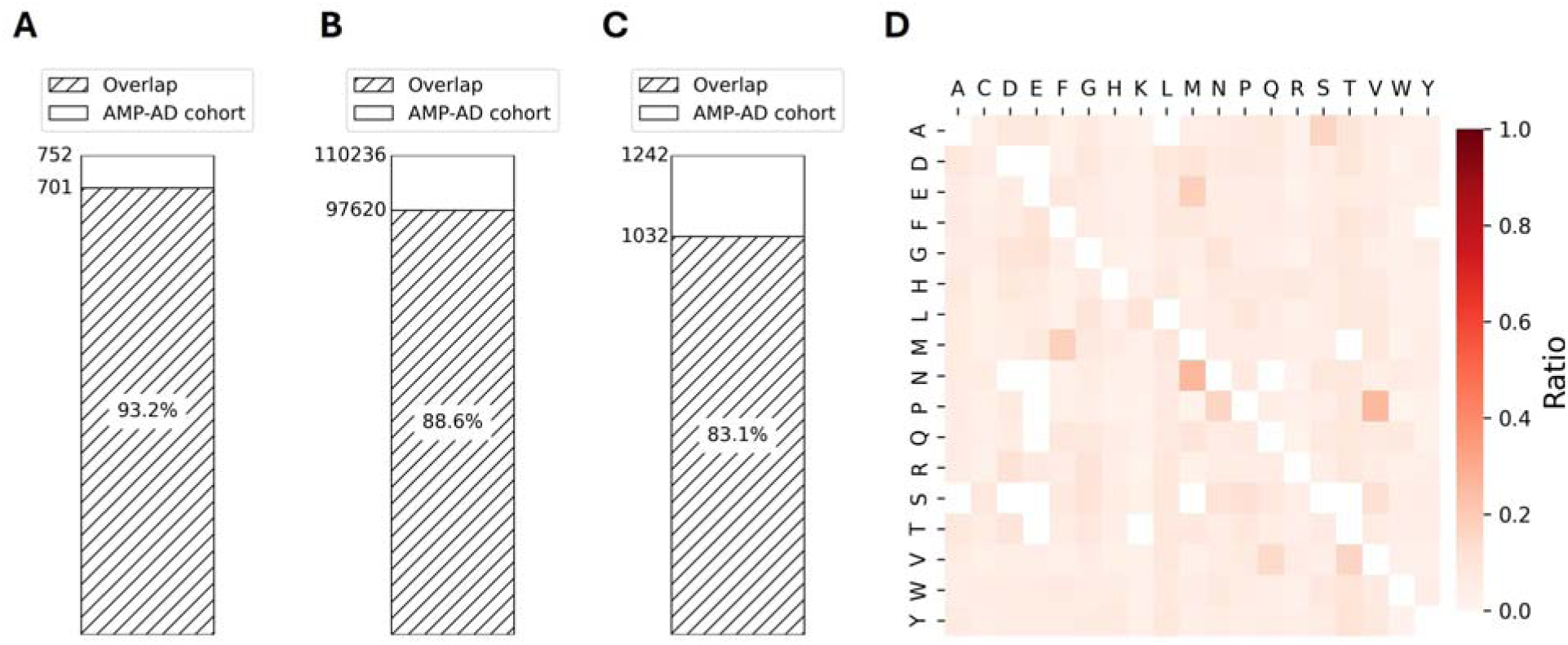
Generalizability of the pipeline in the additional proteomic dataset. (A) The overlap results of identified protein numbers. Slashed section: proteins that are replicated in the additional dataset. (B) The overlap results of PSM numbers. Slashed section: PSMs with the same peptide sequence in the additional dataset. (C) The overlap results of peptide-sequence numbers for the significantly up-regulated UPSPs. Slashed section: peptide sequences that are replicated in the additional dataset. (D) Heatmaps display the normalized ratios for all categories of AA substitutions in the additional dataset. The vertical axis represents the original AAs, while the horizontal axis denotes the AAs following substitution; each row is normalized to sum to one.

Figure 4A illustrates the overlapping proteins detected in the AMP-AD and the additional dataset. In total, 701 out of the 752 proteins that carry a quantified UPSP in the AMP-AD dataset were also identified in the additional dataset, corresponding to an overlap ratio of 93.2%. We next calculated the PSM with the same peptide sequence overlap between the two datasets (Figure 4B), where 97,620 of 110,236 PSMs overlapped, yielding an overlap rate of 88.6%. Furthermore, we evaluated the overlap at the peptide level. The significantly up-regulated UPSPs of the AMP-AD dataset correspond to 1,242 distinct peptide sequences, of which 1,032 were also identified in the additional dataset (Figure 4C), yielding an overlap ratio of 83.1%. Because these three comparisons are matched on protein identity or on the unmodified peptide sequence, they establish that the same proteins and peptides are re-identified in an independent acquisition; replication of the substitution events themselves is assessed at the site level below. Especially, high replication rates are observed for AA substitution sites across the aforementioned key proteins in the additional dataset: 57% for GAPDH (12/21), 80% for ENO1 (12/15), 55% for GFAP (6/11), 90% for ACTG1 (27/30), 86% for HBB (19/22), and 93% for TUBA1B (26/28). These overlap ratios reflect qualitative recurrence of identified substitutions across two distinct proteomic preparations. Besides, the normalized ratio for each type of AA substitution across the additional dataset is represented as heatmaps in Figure 4D, displaying a similar pattern of AA substitutions that aligns with the findings from the AMP-AD dataset. Consistent with the AMP-AD dataset, among all N>X substitutions, the N>M substitution is identified as the most prevalent, with an approximate ratio of 26%. (Supplementary File 5). The search outcomes from the additional dataset have been stored in Zenodo [28] with record number 15753438.

In summary, the substantial overlap ratios observed between the AMP-AD and additional datasets indicate that the substitution events reported here are qualitatively reproducible in an independently acquired AD proteome. Because PXD013753 comprises a single TMT experiment in which 33 individuals were pooled into eight group-level channels, this comparison supports reproducibility across two different proteomic preparations.

## 3 METHODS

### 3.1 AD proteomic dataset and data preprocessing

For this study, we accessed proteomic data from the AMP-AD via the AD Knowledge Portal (https://www.synapse.org/) [22]. The dataset included samples from the STG, contributed by 244 individuals across control and pathologically defined AD cases [23]. To validate the generalizability of our findings, we further retrieved an independent validation dataset from ProteomeXchange under the identifier PXD013753 [76], which included samples from 33 individuals. MS2 spectra that have valid contents were selected and converted to MGF format using MSConvert [77]. We retrieved 1,824 raw files from the AMP-AD repository. However, during data preprocessing, one file was excluded due to missing valid MS2 spectra in the resulting MGF file. Consequently, subsequent analyses utilized a total of 1,823 MGF files.

### 3.2 AD proteomic data analysis

The FASTA file for human proteins was retrieved from UniProt [29]. We set the following parameters for PIPI-C: an MS1 tolerance of 10 ppm, an MS2 tolerance of 0.01 Da, trypsin as the enzyme allowing up to two missed cleavages, and a peptide length range of 6-50 AAs. Fixed modifications included carbamidomethylation (C, +57.0215 Da) and the isobaric label on lysine. Because the two datasets were labelled with different reagents, this fixed modification was TMTpro (K, +304.2071 Da) for AMP-AD and TMT (K, +229.1629 Da) for PXD013753. Variable modifications encompassed the same reagent on S and the peptide N-terminus (TMTpro for AMP-AD, TMT for PXD013753) and all possible single AA substitutions (using UNIMOD [24]), with a [-250 Da, +250 Da] modification search range. Isoleucine and leucine were regarded as equivalent in this study. Decoy proteins were incorporated to manage the false discovery rate (FDR) using a target-decoy approach by appending shuffled decoy sequences to the target FASTA, while all other parameters were maintained at their default settings.

In the Comet search, we re-searched only those spectra whose PIPI-C identification carried at least one AA substitution and whose substitution-bearing peptidoform had a complete PIPI-C modification profile shared by at least two spectra within the same raw file. For each such spectrum, the per-spectrum modification profile reported by PIPI-C was supplied to Comet as the variable-modification list. The variable modifications applied in Comet were directly based on the PIPI-C results corresponding to each specific spectrum. All other parameters for the Comet search were consistent with those utilized in PIPI-C. Only PSMs that were identified by both search engines were preserved for bioinformatic analysis. This re-scoring step does not constitute independent validation in the strict sense, since the variable-modification list per spectrum is inherited from PIPI-C; rather, it filters out identifications that fail to reproduce under a different scoring function.

### 3.3 Quantification details

Quantification was restricted to PSMs whose TMTpro labeling pattern was consistent with the experimental design. Peptides supported by multiple PSMs were classified as quantifiable. Reporter ion intensities corresponding to each PSM were extracted directly from the MS2. We assumed that all samples contained unregulated peptides with abundances akin to those found in the common reference (CR) sample. Consequently, after normalization, these peptides were expected to yield log-ratios centered at zero. To achieve this, we implemented a two-component normalization of TMT label intensities using CR intensities as the reference standard [23].

At the PSM level, log ratios were calculated as the average of log-transformed values across AD patients for each PSM. Peptides with the same modification profiles were combined into UPSP. The UPSP-level ratio was calculated by averaging all constituent PSM-level TMT ratios within each subgroup. Then, we derived experiment-wide average log ratios for each UPSP. For downstream analysis, we retained only UPSPs detected in more than 60% of the 19 TMTpro experiments (i.e., in at least 12 of 19 experiments) for quantification. This criterion applies to the AMP-AD dataset only; PXD013753 consists of a single TMT experiment and is therefore not subject to it. For each retained UPSP, statistical significance of the deviation of its experiment-wide average log ratio from zero was assessed using one-sample t-tests across experiments, with Benjamini-Hochberg FDR correction applied to generate adjusted p-values for each UPSP. Because all ratios are computed against the CR channel, an up-regulated UPSP throughout this work denotes a UPSP whose abundance in AD samples is significantly higher than that in the common reference, rather than one showing a significant difference between AD and control samples.

The case-control contrast reported in Section 2.2 was computed separately. AD and control status follows the harmonized neuropathological classification distributed with the dataset. Reporter intensities were first divided by the total reporter signal of their channel within each experiment, and log ratios were taken against the mean of that experiment’s global internal standard channels. Within each experiment the AD channels and the control channels were averaged and their difference taken, giving one paired estimate per peptidoform per experiment; because the two contributing institutions occupy systematically different reporter channels, this was done separately within Emory and within Mayo Clinic channels. The per-institution mean and standard error across experiments were combined by inverse-variance fixed-effect meta-analysis and referred to a t distribution, with Benjamini-Hochberg correction applied once across the 1,678 substitution-bearing peptidoforms testable in both institutions, and events were retained only when the effect was positive in both. Between-institution heterogeneity is reported for every event in Supplementary File 6.

### 3.4 Substitution occupancy

Adapting a biotin occupancy-like metric [26, 27], the N>X substitution occupancy of a protein was defined as the number of asparagine residues at which an N>X substitution was detected, divided by the asparagine density of that protein, where the asparagine density is the number of asparagine residues in the protein divided by the protein length; N>M occupancy was defined analogously using only N>M events. This normalization prevents proteins that are simply asparagine-rich from dominating the ranking. Occupancy was computed only for proteins identified in more than 60% of experiments.

### 3.5 GO enrichment analysis

We performed GO enrichment analysis on proteins containing quantified UPSPs that met the following criteria: (1) q-value < 0.05 in quantification results, and (2) the presence of AA substitution modifications. The set of proteins identified in this dataset served as the reference background for these analyses, rather than the whole human proteome, so that categories which are intrinsically over-represented among proteins accessible to shotgun proteomics are not spuriously enriched. The background comprised the 3,389 leading proteins of all Comet-validated PSMs, of which 3,371 carried a Gene Ontology annotation and entered the test. The study set for this analysis was defined before the exclusion of ambiguous mass shifts listed in Table 1 and therefore comprises 187 leading proteins, of which 185 carried an annotation in the detected-proteome background. The complete results under both backgrounds are provided as Supplementary File 7. The statistical significance of the enrichment was evaluated using the two-tailed Fisher’s exact test, incorporating BH FDR correction (q-value < 0.05). All GO enrichment analyses were implemented using GOATOOLS [78] in Python.

### 3.6 Details of protein-protein interaction network

To investigate functional protein networks, we obtained PPI data from STRING [79], which includes interactions from multiple evidence channels, including experimental data, curated databases, text mining, co-expression, genomic context, and phylogenetic profiling. The PPI network was constructed and visualized using Cytoscape (v3.10.2) [80], with proteins represented as nodes and their interactions as edges.

### 3.7 Details of protein structure prediction and comparison

Alignment and modeling of protein structures were performed using ChimeraX. The structure of GAPDH was obtained through cryo-electron microscopy (PDB: 8G17). The crystal structure of ENO1, derived from human sources and expressed in Escherichia coli, was used as a reference (PDB: 2PSN). The structure of human non-muscle gamma-actin (ACTG1) was acquired via cryo-electron microscopy at 3.4 A resolution (PDB: 8DNF), while the cryo-electron microscopy structure of human TUBA1B was sourced from cabbage looper expression (PDB: 5IJ0). For variant structure prediction, we submitted protein sequences with cumulative AA substitutions to AlphaFold3’s web service, incorporating only the most strongly regulated AA substitution (highest regulation ratio) at each variable site. We acknowledge that cumulative substitution modeling does not represent any single physical proteoform observed in the data; the resulting structures are interpreted as spatial hotspot maps rather than as functional structural predictions. Among the generated structures, we selected the prediction with the highest pLDDT score for structural comparisons.

## 4 DISCUSSION

This study performed AA substitutomic analysis using mass spectrometry-based proteomic data from the AMP-AD Consortium, which includes samples from the STG of 244 individuals across control and pathologically defined AD cases. Our application of the substitutomics framework [19] to AD proteomic data provides the first proteome-wide map of AA substitutions in AD. To optimize proteomic analysis, we used a mixed search pipeline that maximizes both proteome coverage and identification precision. Using PIPI-C [20], our method comprehensively detects all possible AA substitutions without restricting the number of substitutions per PSM. Considering the risk of false-positive identifications in PIPI-C results, we re-scored the substitution-bearing spectra with Comet [21], which removes identifications that do not reproduce under a different scoring function. The combination of these two search strategies achieves a robust balance between comprehensive proteomic exploration and high-confidence identification.

We observed consistent AA substitution patterns across both the AMP-AD dataset and the dataset of PXD013753, with the N>M substitution representing the most frequent event among all N>X substitutions. This cross-dataset consistency yields novel mechanistic insight into AA substitutions in AD research: it confirms that the observed N>M substitution is a disease-associated molecular signature rather than random stochastic variation, a distinction of high significance for distinguishing biologically meaningful alterations from background translational noise. It is known that translation fidelity is a critical factor influencing gene expression and protein function, and ribosomal errors during mRNA translation occur at an estimated stochastic rate of 10^−4^ per codon [81]. Based on these observations, we propose that the high frequency of the N>M substitution may arise from two interrelated factors: (1) enhanced metabolic dependence on methionine in AD cells, a phenomenon documented in previous cancer studies [82]; and (2) dysfunction of methionyl-tRNA in the nervous systems of AD patients [83]. The consistent association of this high-frequency substitution with AD indicates its potential as a promising therapeutic target.

AA substitutions play critical roles in cellular processes and biological functions and arise from three major sources: (1) DNA mutations, (2) mRNA mutations, and (3) errors during protein translation. In this study, we employ the AA substitutomic analysis pipeline to establish a high-throughput method for investigating proteome-level AA substitutions. Using this pipeline, we demonstrate that AA substitutions not recorded in genomic or transcriptomic variant databases are prevalent in AD patient cells. Furthermore, we identify literature-reported substitution events occurring in AD-associated proteins, which can alter protein structural conformation and biological function. This study provides a novel, protein-centric perspective to understand AD pathogenesis, offers promising molecular candidates for AD biomarker screening, and identifies candidates for follow-up.

While previous work has focused almost exclusively on genetically or transcriptionally encoded substitutions, proteomic research has long been limited by the absence of high-throughput open-search pipelines for unbiased, large-scale AA substitution identification. To address this bottleneck, we applied a robust AA substitutomics pipeline for the high-throughput detection of protein-level substitutions, enabling quantitative proteomic analysis of their regulatory consequences. We validated our results by cross-referencing significantly regulated substitutions with known variants in the UniProt database [29]. A 5.4% overlap with documented UniProt variants (42 of 784 substitution events; Supplementary File 4) corroborates our observations. Since UniProt-curated substitutions are mostly derived from genomic or transcriptomic mutations, the majority of the substitutions identified here are not attributable to any catalogued germline or somatic variant; their origin, however, cannot be assigned from the MS data alone and requires orthogonal validation. The same pipeline is applicable to any isobaric-labelled dataset and is not specific to AD.

This study identified multiple significantly regulated AA substitutions in key AD-associated proteins, highlighting their potential biological relevance to AD progression. Nevertheless, several important limitations of this study should be noted. First, the vast combinatorial complexity of AA substitutions, where any residue can theoretically be replaced by 19 other AAs, creates substantial challenges for mechanistic interpretation. Current functional annotations remain insufficient to fully distinguish causal mechanisms from such complex substitution patterns.

The reference-relative quantification establishes which substitution events are reproducibly detected across experiments, whereas the institution-stratified case-control analysis in Section 2.2 establishes which of them differ between AD and neuropathological controls; the two answer different questions. The latter resolves a concentrated rather than a comprehensive set, 138 events in 52 proteins with half of them in five proteins, and it is limited by the number of TMTpro experiments available within each institution rather than by the quality of the identifications; consistent with the institution-associated variance previously documented for this dataset [23], larger cohorts matched within institution would be expected to resolve more of them. In addition, technical constraints of mass spectrometry affect our interpretations: the method captures population-averaged proteomic profiles rather than single-molecule resolution. Thus, the detected substitutions likely occur across different protein molecules rather than simultaneously on the same protein. Our structural modeling reveals their cumulative effects but cannot determine their precise co-occurrence. Finally, the limited availability of clinical samples and experimental restrictions hinder immediate large-scale wet-lab validation. Therefore, fully elucidating the role of AA substitutomics in AD will require interdisciplinary collaborative efforts to bridge computational predictions with mechanistic experimental validation.

## 5 CONCLUSION

AA substitutions serve a pivotal role in regulating cellular processes in AD, mediating dysregulated signaling pathways and cellular functions. Previous studies have been restricted by technical limitations in peptide identification, which confined analyses to a narrow range of potential substitution events. To overcome this critical bottleneck in AD research, we employed PIPI-C [20], a state-of-the-art open-search tool that enables unbiased and comprehensive profiling of all possible

AA substitutions. Among the AA substitutions and their respective locations identified, 5.4% overlap with UniProt-curated germline/somatic variants; the remaining 94.6% are not catalogued in UniProt, indicating events accessible primarily through proteomic profiling, though their molecular origin (post-translational or unannotated genetic) cannot be resolved from MS data alone.

This study reports a comprehensive AA substitutomics analysis of AD-related proteins independent of genomic or transcriptomic data. Key findings include the identification of two clinically significant substitution sites (242 and 352) in GFAP, which are previously linked to Alexander disease, a rare neurodegenerative disorder. In ACTG1, we detected a substitution at residue 370 (V370R), the position at which the reported V370A mutation impairs cell proliferation under thermal and hyperosmotic stress. Additionally, we detected three well-documented substitution sites (111, 115, and 116) in HBB that trigger a severe thalassemia-like phenotype via hemoglobin dysfunction. While a subset of the identified substitutions is supported by existing literature, the majority represent novel discoveries that require further experimental validation. The limited mechanistic understanding of these unreported substitutions emphasizes the urgent need for large-scale proteomic studies to elucidate the specific functions of AA substitutions in AD pathogenesis.

In conclusion, this study addresses the technical bottleneck in large-scale profiling of multiple AA substitutions within single peptides and establishes a high-throughput, robust workflow for the systematic characterization of AA substitutions, especially those that are not catalogued in genomic or transcriptomic variant databases and therefore escape traditional genomic and transcriptomic screens. Our customized AA substitutomic pipeline reveals the widespread prevalence of proteome-level AA substitutions in AD samples and maps these substitution events to core AD-associated proteins, further demonstrating their functional consequences, such as impaired protein stability and dysregulated protein–protein interactions. Validated by high consistency with published experimental evidence, our proteome-wide pipeline enables the first comprehensive landscape of AA substitutions in AD. Collectively, our findings highlight AA substitutions as critical modulators of protein function that potentially drive AD pathogenesis, offer a perspective for understanding neurodegeneration, and lay a foundation for developing substitution-oriented diagnostic biomarkers and targeted therapeutic strategies.

## 6 DATA AVAILABILITY AND SUPPLEMENTARY FILES

The results of the AMP-AD proteomic dataset are available on Zenodo, with the record number 15010408. https://zenodo.org/records/15010408

The results of the additional proteomic dataset are available on Zenodo, with the record number 15753438. https://zenodo.org/records/15753438

Supplementary File 1: The normalized ratio of each AA substitution in the AMP-AD dataset.

Supplementary File 2: The N>M and N>X substitution occupancy in the AMP-AD dataset.

Supplementary File 3: The quantification results of the AMP-AD dataset. It contains the AA substitutions, peptide sequence, regulation ratio, and statistics of the results.

Supplementary File 4: The AA substitution details of all proteins in the AMP-AD dataset, and the AA substitutions that have been recorded in the UniProt database.

Supplementary File 5: The normalized ratio of each AA substitution in the additional AD dataset.

Supplementary File 6: The 138 substitution-bearing peptidoforms that are significantly more abundant in AD than in neuropathological controls in both contributing institutions, with the meta-analysed effect size and adjusted p-value and the underlying estimates obtained within Emory and within Mayo Clinic.

Supplementary File 7: The complete GO enrichment results for the substitution-bearing proteins under both the whole-genome and the identified-proteome background.

## 7 ACKNOWLEDGEMENTS

We thank the Generic Diagramming Platform (https://biogdp.com/) for offering the schematic diagram.

## 8 Declarations

### Funding

This work was partially supported by ITC grants MHP/033/20 and ITS/043/23, RGC grants 16102422, 16103621, R4012-18, C7015-23G, and T12-101/23N from the Hong Kong SAR government, Hetao Cooperation Zone Project HzQB-KCZYB-2020083, and internal grants 3030_009, BGF.001.2023, CSSET24SC01, Z1056, and OKT26EG06 from HKUST.

### Conflicts of interest/Competing interests

The authors declare no competing interests.

### Ethics approval and consent to participate

This study is a secondary computational analysis of previously published, de-identified proteomic datasets (the AMP-AD consortium dataset [22, 23] and the ProteomeXchange dataset PXD013753 [76]). No new human participants, human tissue, or human data were collected by the authors. Ethics approval and informed consent for sample collection were obtained in the respective original studies. Therefore, no additional ethics approval was required for this study.

### Consent for publication

Not applicable. This study did not include any individual person’s identifiable data, images, or clinical details requiring consent for publication.

### Availability of data and material

The results of the AMP-AD proteomic dataset are available on Zenodo (record 15010408, https://zenodo.org/records/15010408). The results of the additional proteomic dataset are available on Zenodo (record 15753438, https://zenodo.org/records/15753438). The original datasets analyzed in this study are publicly available: the AMP-AD dataset from the AMP-AD consortium [22, 23] and dataset PXD013753 from ProteomeXchange [76]. Supplementary Files 1–7 and the Supplementary Note are provided with this article.

### Authors’ contributions

P.Z., S.L., and S.D. designed the study and developed the analysis pipeline. P.Z. and S.L. performed the data analysis. S.D. and C.Z. carried out the biological interpretation and validation. P.Z. wrote the original draft. N.L. and W.Y. supervised the project, acquired funding, and revised the manuscript. All authors read and approved the final manuscript.

## Notes

### Competing Interest Statement

The authors have declared no competing interest.

## References

[1] L. X. Tay, S. C. Ong, L. J. Tay, T. Ng, and T. Parumasivam. “Economic burden of Alzheimer’s disease: A systematic review”. Value in Health Regional Issues, 40, 1–12, 2024.

[2] P. Scheltens, K. Blennow, M. M. Breteler, B. De Strooper, G. B. Frisoni, S. Salloway, and W. M. Van der Flier. “Alzheimer’s disease”. The Lancet, 388(10043), 505–517, 2016.

[3] C. Haass, A. Y. Hung, D. J. Selkoe, and D. B. Teplow. “Mutations associated with a locus for familial Alzheimer’s disease result in alternative processing of amyloid beta-protein precursor”. Journal of Biological Chemistry, 269(26), 17741–17748, 1994.

[4] M. Seuma, B. Lehner, and B. Bolognesi. “An atlas of amyloid aggregation: The impact of substitutions, insertions, deletions and truncations on amyloid beta fibril nucleation”. Nature Communications, 13(1), 7084, 2022.

[5] L. Connelly, H. Jang, F. Teran Arce, S. Ramachandran, B. L. Kagan, R. Nussinov, and R. Lal. “Effects of point substitutions on the structure of toxic Alzheimer’s *β*-amyloid channels: Atomic force microscopy and molecular dynamics simulations”. Biochemistry, 51(14), 3031– 3038, 2012.

[6] M. D. Topal and J. R. Fresco. “Complementary base pairing and the origin of substitution mutations”. Nature, 263(5575), 285–289, 1976.

[7] S. Seshadri, A. L. Fitzpatrick, M. A. Ikram, A. L. DeStefano, V. Gudnason, M. Boada, J. C. Bis, A. V. Smith, M. M. Carrasquillo, J. C. Lambert, et al. “Genome-wide analysis of genetic loci associated with Alzheimer disease”. JAMA, 303(18), 1832–1840, 2010.

[8] L. Bertram and R. E. Tanzi. “Genome-wide association studies in Alzheimer’s disease”. Human Molecular Genetics, 18(R2), R137–R145, 2009.

[9] R. N. Rosenberg, D. Lambracht Washington, G. Yu, and W. Xia. “Genomics of Alzheimer disease: A review”. JAMA Neurology, 73(7), 867–874, 2016.

[10] H. Mathys, J. Davila Velderrain, Z. Peng, F. Gao, S. Mohammadi, J. Z. Young, M. Menon, L. He, F. Abdurrob, X. Jiang, et al. “Single-cell transcriptomic analysis of Alzheimer’s disease”. Nature, 570(7761), 332–337, 2019.

[11] H. Patel, A. K. Hodges, C. Curtis, S. H. Lee, C. Troakes, R. J. Dobson, and S. J. Newhouse. “Transcriptomic analysis of probable asymptomatic and symptomatic alzheimer brains”. Brain, Behavior, and Immunity, 80, 644–656, 2019.

[12] H. Patel, R. J. Dobson, and S. J. Newhouse. “A meta-analysis of Alzheimer’s disease brain transcriptomic data”. Journal of Alzheimer’s Disease, 68(4), 1635–1656, 2019.

[13] S. Dujardin, C. Commins, A. Lathuiliere, P. Beerepoot, A. R. Fernandes, T. V. Kamath, M. B. De Los Santos, N. Klickstein, D. L. Corjuc, B. T. Corjuc, et al. “Tau molecular diversity contributes to clinical heterogeneity in Alzheimer’s disease”. Nature Medicine, 26(8), 1256– 1263, 2020.

[14] M. H. Abreha, E. B. Dammer, L. Ping, T. Zhang, D. M. Duong, M. Gearing, J. J. Lah, A. I. Levey, and N. T. Seyfried. “Quantitative analysis of the brain ubiquitylome in Alzheimer’s disease”. Proteomics, 18(20), 1800108, 2018.

[15] E. B. Dammer, C. H. Na, P. Xu, N. T. Seyfried, D. M. Duong, D. Cheng, M. Gearing, H. Rees, J. J. Lah, A. I. Levey, et al. “Polyubiquitin linkage profiles in three models of proteolytic stress suggest the etiology of Alzheimer disease”. Journal of Biological Chemistry, 286(12), 10457–10465, 2011.

[16] Q. Zhang, C. Ma, L. S. Chin, and L. Li. “Integrative glycoproteomics reveals protein Nglycosylation aberrations and glycoproteomic network alterations in Alzheimer’s disease”. Science Advances, 6(40), eabc5802, 2020.

[17] Y. Cai, S. S. A. An, and S. Kim. “Mutations in presenilin 2 and its implications in Alzheimer’s disease and other dementia-associated disorders”. Clinical Interventions in Aging, 1163–1172, 2015.

[18] R. J. Kelleher III and J. Shen. “Presenilin-1 mutations and Alzheimer’s disease”. Proceedings of the National Academy of Sciences, 114(4), 629–631, 2017.

[19] P. Zhao, S. Dai, S. Lai, C. Zhou, N. Li, and W. Yu. “Amino acid substitutomics: Profiling amino acid substitutions at proteomic scale unveils biological implication and escape mechanism in cancer”. bioRxiv, 2024–11, 2026.

[20] S. Lai, S. Dai, P. Zhao, C. Zhou, N. Li, and W. Yu. “PIPI-C: A combinatorial optimization framework for identifying post-translational modification hot-spots in mass spectrometry data”. Molecular & Cellular Proteomics, 25(2), 101494, 2026.

[21] J. K. Eng, T. A. Jahan, and M. R. Hoopmann. “Comet: An open-source MS/MS sequence database search tool”. Proteomics, 13(1), 22–24, 2013.

[22] R. J. Hodes and N. Buckholtz. “Accelerating medicines partnership: Alzheimer’s disease (AMP-AD) knowledge portal aids Alzheimer’s drug discovery through open data sharing”. Expert Opinion on Therapeutic Targets, 20(4), 389–391, 2016.

[23] F. Seifar, E. J. Fox, A. Shantaraman, Y. Liu, E. B. Dammer, E. Modeste, D. M. Duong, L. Yin, A. N. Trautwig, Q. Guo, et al. “Large-scale deep proteomic analysis in Alzheimer’s disease brain regions across race and ethnicity”. bioRxiv, 2024.

[24] D. M. Creasy and J. S. Cottrell. “Unimod: Protein modifications for mass spectrometry”. Proteomics, 4(6), 1534–1536, 2004.

[25] X. Gabaix. “Zipf’s law for cities: An explanation”. The Quarterly Journal of Economics, 114(3), 739–767, 1999.

[26] N. Yang, J. Ren, S. Dai, K. Wang, M. Leung, Y. Lu, Y. An, A. Burlingame, S. Xu, Z. Wang, et al. “The quantitative biotinylproteomics studies reveal a wind-related kinase 1 (Raf-Like kinase 36) functioning as an early signaling component in wind-induced thigmomorphogenesis and gravitropism”. Molecular & Cellular Proteomics, 23(3), 100738, 2024.

[27] K. Wu, N. Yang, J. Ren, S. Liu, K. Wang, S. Dai, Y. Lu, Y. An, F. Tian, Z. Gao, et al. “Cytosolic WPRa4 and plastoskeletal PMI4 proteins mediate touch response in a model organism arabidopsis”. Molecular & Cellular Proteomics, 24(7), 101015, 2025.

[28] European Organization For Nuclear Research and OpenAIRE. Zenodo. en. doi: 10.25495/7GXK-RD71. url: https://www.zenodo.org/.

[29] U. Consortium. “UniProt: a worldwide hub of protein knowledge”. Nucleic Acids Research, 47(D1), D506–D515, 2019.

[30] M. Ashburner, C. A. Ball, J. A. Blake, D. Botstein, H. Butler, J. M. Cherry, A. P. Davis, K. Dolinski, S. S. Dwight, J. T. Eppig, et al. “Gene ontology: Tool for the unification of biology”. Nature Genetics, 25(1), 25–29, 2000.

[31] A. Smith, S. P. Datta, G. H. Smith, P. N. Campbell, R. Bentley, and H. A. McKenzie. Oxford dictionary of biochemistry and molecular biology. Revised edition.

[32] A. S. Verkman. “Solute and macromolecule diffusion in cellular aqueous compartments”. Trends in Biochemical Sciences, 27(1), 27–33, 2002.

[33] L. E. Goldstein, J. A. Muffat, R. A. Cherny, R. D. Moir, M. H. Ericsson, X. Huang, C. Mavros, J. A. Coccia, K. Y. Faget, K. A. Fitch, et al. “Cytosolic *β*-amyloid deposition and supranuclear cataracts in lenses from people with Alzheimer’s disease”. The Lancet, 361(9365), 1258–1265, 2003.

[34] K. Iqbal, A. d. C. Alonso, and I. Grundke Iqbal. “Cytosolic abnormally hyperphosphorylated tau but not paired helical filaments sequester normal MAPs and inhibit microtubule assembly”. Journal of Alzheimer’s Disease, 14(4), 365–370, 2008.

[35] J. Hardy and D. J. Selkoe. “The amyloid hypothesis of Alzheimer’s disease: progress and problems on the road to therapeutics”. Science, 297(5580), 353–356, 2002.

[36] E. Karran, M. Mercken, and B. D. Strooper. “The amyloid cascade hypothesis for Alzheimer’s disease: An appraisal for the development of therapeutics”. Nature Reviews Drug Discovery, 10(9), 698–712, 2011.

[37] K. Wakasugi, T. Nakano, and I. Morishima. “Oxidative stress-responsive intracellular regulation specific for the angiostatic form of human tryptophanyl-tRNA synthetase”. Biochemistry, 44(1), 225–232, 2005.

[38] Y. Song, Q. Luo, H. Long, Z. Hu, T. Que, X. Zhang, Z. Li, G. Wang, L. Yi, Z. Liu, et al. “Alpha-enolase as a potential cancer prognostic marker promotes cell growth, migration, and invasion in glioma”. Molecular Cancer, 13, 1–12, 2014.

[39] L. Shen, C. Chen, A. Yang, Y. Chen, Q. Liu, and J. Ni. “Redox proteomics identification of specifically carbonylated proteins in the hippocampi of triple transgenic Alzheimer’s disease mice at its earliest pathological stage”. Journal of Proteomics, 123, 101–113, 2015.

[40] R. Sultana, R. A. Robinson, F. Di Domenico, H. M. Abdul, D. K. S. Clair, W. R. Markesbery, J. Cai, W. M. Pierce, and D. A. Butterfield. “Proteomic identification of specifically carbonylated brain proteins in APPNLh/APPNLh × PS-1P264L/PS-1P264L human double mutant knock-in mice model of Alzheimer disease as a function of age”. Journal of Proteomics, 74(11), 2430–2440, 2011.

[41] W. O. Opii, G. Joshi, E. Head, N. W. Milgram, B. A. Muggenburg, J. B. Klein, W. M. Pierce, C. W. Cotman, and D. A. Butterfield. “Proteomic identification of brain proteins in the canine model of human aging following a long-term treatment with antioxidants and a program of behavioral enrichment: Relevance to Alzheimer’s disease”. Neurobiology of Aging, 29(1), 51– 70, 2008.

[42] I. Ferrer, R. Blanco, M. Carmona, and B. Puig. “Phosphorylated c-MYC expression in Alzheimer disease, Pick’s disease, progressive supranuclear palsy and corticobasal degeneration”. Neuropathology and Applied Neurobiology, 27(5), 343–351, 2001.

[43] H. g. Lee, G. Casadesus, A. Nunomura, X. Zhu, R. J. Castellani, S. L. Richardson, G. Perry, D. W. Felsher, R. B. Petersen, and M. A. Smith. “The neuronal expression of MYC causes a neurodegenerative phenotype in a novel transgenic mouse”. The American Journal of Pathology, 174(3), 891–897, 2009.

[44] D. K. Cullen, C. M. Simon, and M. C. LaPlaca. “Strain rate-dependent induction of reactive astrogliosis and cell death in three-dimensional neuronal-astrocytic co-cultures”. Brain Research, 1158, 103–115, 2007.

[45] K. Y. Kim, K. Y. Shin, and K. A. Chang. “GFAP as a potential biomarker for Alzheimer’s disease: A systematic review and meta-analysis”. Cells, 12(9), 1309, 2023.

[46] J. B. Pereira, S. Janelidze, R. Smith, N. Mattsson Carlgren, S. Palmqvist, C. E. Teunissen, H. Zetterberg, E. Stomrud, N. J. Ashton, K. Blennow, et al. “Plasma GFAP is an early marker of amyloid-*β* but not tau pathology in Alzheimer’s disease”. Brain, 144(11), 3505–3516, 2021.

[47] J. Gorospe, S. Naidu, A. Johnson, V. Puri, G. Raymond, S. Jenkins, R. Pedersen, D. Lewis, P. Knowles, R. Fernandez, et al. “Molecular findings in symptomatic and pre-symptomatic Alexander disease patients”. Neurology, 58(10), 1494–1500, 2002.

[48] R. Li, A. B. Johnson, G. Salomons, J. E. Goldman, S. Naidu, R. Quinlan, B. Cree, S. Z. Ruyle, B. Banwell, M. D’Hooghe, et al. “Glial fibrillary acidic protein mutations in infantile, juvenile, and adult forms of Alexander disease”. Annals of Neurology, 57(3), 310–326, 2005.

[49] W. Choi, H. Wu, K. Yserentant, B. Huang, and Y. Cheng. “Efficient tagging of endogenous proteins in human cell lines for structural studies by single-particle cryo-EM”. Proceedings of the National Academy of Sciences, 120(31), e2302471120, 2023.

[50] H. J. Kang, S. K. Jung, S. J. Kim, and S. J. Chung. “Structure of human *α*-enolase (hENO1), a multifunctional glycolytic enzyme”. Biological Crystallography, 64(6), 651–657, 2008.

[51] E. M. Mandelkow, J. Biernat, G. Drewes, N. Gustke, B. Trinczek, and E. Mandelkow. “Tau domains, phosphorylation, and interactions with microtubules”. Neurobiology of Aging, 16(3), 355–362, 1995.

[52] J. Avila. “Tau kinases and phosphatases”. Journal of Cellular and Molecular Medicine, 12(1), 258, 2008.

[53] D. H. Lau, M. Hogseth, E. C. Phillips, M. J. O’Neill, A. M. Pooler, W. Noble, and D. P. Hanger. “Critical residues involved in tau binding to fyn: implications for tau phosphorylation in Alzheimer’s disease”. Acta Neuropathologica Communications, 4, 1–13, 2016.

[54] W. Gulisano, D. Maugeri, M. A. Baltrons, M. F’a, A. Amato, A. Palmeri, L. D’Adamio, C. Grassi, D. Devanand, L. S. Honig, et al. “Role of amyloid-*β* and tau proteins in Alzheimer’s disease: confuting the amyloid cascade”. Journal of Alzheimer’s Disease, 64(s1), S611–S631, 2018.

[55] H. O. Villar and L. M. Kauvar. “Amino acid preferences at protein binding sites”. FEBS Letters, 349(1), 125–130, 1994.

[56] F. Ceppi, C. Langlois Pelletier, V. Gagne, J. Rousseau, C. Ciolino, S. D. Lorenzo, K. M. Kevin, D. Cijov, S. E. Sallan, L. B. Silverman, et al. “Polymorphisms of the vincristine pathway and response to treatment in children with childhood acute lymphoblastic leukemia”. Pharmacogenomics, 15(8), 1105–1116, 2014.

[57] M. Li, R. Geng, C. Li, F. Meng, H. Zhao, J. Liu, J. Dai, and X. Wang. “Dysregulated gene-associated biomarkers for Alzheimer’s disease and aging”. Translational Neuroscience, 12(1), 83–95, 2021.

[58] N. D. Rendtorff, M. Zhu, T. Fagerheim, T. L. Antal, M. Jones, T. M. Teslovich, E. M. Gillanders, M. Barmada, E. Teig, J. M. Trent, et al. “A novel missense mutation in ACTG1 causes dominant deafness in a Norwegian DFNA20/26 family, but ACTG1 mutations are not frequent among families with hereditary hearing impairment”. European Journal of Human Genetics, 14(10), 1097–1105, 2006.

[59] L. R. Otterbein, C. Cosio, P. Graceffa, and R. Dominguez. “Crystal structures of the vitamin D-binding protein and its complex with actin: Structural basis of the actin-scavenger system”. Proceedings of the National Academy of Sciences, 99(12), 8003–8008, 2002.

[60] V. A. Klenchin, J. S. Allingham, R. King, J. Tanaka, G. Marriott, and I. Rayment. “Trisoxazole macrolide toxins mimic the binding of actin-capping proteins to actin”. Nature Structural & Molecular Biology, 10(12), 1058–1063, 2003.

[61] M. Hertzog, C. Van Heijenoort, D. Didry, M. Gaudier, J. Coutant, B. Gigant, G. Didelot, T. Preat, M. Knossow, E. Guittet, et al. “The *β*-thymosin/WH2 domain: structural basis for the switch from inhibition to promotion of actin assembly”. Cell, 117(5), 611–623, 2004.

[62] M. Walser, A. Hansen, P. A. Svensson, M. Jern°as, J. Oscarsson, J. Isgaard, and N. D. °Aberg. “Peripheral administration of bovine GH regulates the expression of cerebrocortical betaglobin, GABAB receptor 1, and the lissencephaly-1 protein (LIS-1) in adult hypophysectomized rats”. Growth Hormone & IGF Research, 21(1), 16–24, 2011.

[63] Y. Kobayashi, Y. Fukumaki, N. Komatsu, Y. Ohba, T. Miyaji, and Y. Miura. “A novel globin structural mutant, Showa-Yakushiji (*β*110 Leu-Pro) causing a *β*-thalassemia phenotype”. Blood, 70(5), 1688–1691, 1987.

[64] S. Murru, D. Poddie, G. Sciarratta, S. Agosti, M. Baffico, C. Melevendi, M. Pirastu, and A. Cao. “A novel *β*-globin structural mutant, Hb Brescia (*β*114 Leu-Pro), causing a severe *β*-thalassemia intermedia phenotype”. Human Mutation, 1(2), 124–128, 1992.

[65] C. M. De Castro, B. Devlin, D. E. Fleenor, M. E. Lee, and R. E. Kaufman. “A novel betaglobin mutation, beta Durham-NC [beta 114 Leu Pro], produces a dominant thalassemia-like phenotype”. Blood, 83(4), 1109–1116, 1994.

[66] V. Divoky, M. Svobodova, K. Indrak, L. Chrobak, T. Molchanova, and T. Huisman. “HB Hradec Králové (HB HK) or *α*2 *β*2 115 (617) Ala Asp, a severely unstable hewglobin variant resulting in a dominant *β*-thalassemia trait in a Czech family”. Hemoglobin, 17(4), 319–328, 1993.

[67] H. J. Kazazian, C. E. Dowling, R. L. Hurwitz, M. Coleman, A. Stopeck, and J. 3. Adams. “Dominant thalassemia-like phenotypes associated with mutations in exon 3 of the beta-globin gene”. Blood, 79(11), 3014–3018, 1992.

[68] E. Domingues-Hamdi, C. Vasseur, J.-B. Fournier, M. C. Marden, H. Wajcman, and V. BaudinCreuza. “Role of *α*-globin H helix in the building of tetrameric human hemoglobin: interaction with *α*-hemoglobin stabilizing protein (AHSP) and heme molecule”. PLoS One, 9(11), e111395, 2014.

[69] D. I. Zafeiriou, M. Economou, and M. Athanasiou-Metaxa. “Neurological complications in *β*-thalassemia”. Brain and Development, 28(8), 477–481, 2006.

[70] Y. G. Chen, T. Y. Lin, H. J. Chen, M. S. Dai, C. L. Ho, and C. H. Kao. “Thalassemia and risk of dementia: A nationwide population-based retrospective cohort study”. European Journal of Internal Medicine, 26(7), 554–559, 2015.

[71] G. Fu, S. Yan, C. J. Khoo, V. C. Chao, Z. Liu, M. Mukhi, R. Hervas, X. D. Li, and S. Ti. “Integrated regulation of tubulin tyrosination and microtubule stability by human *α*-tubulin isotypes”. Cell Reports, 42(6), 2023.

[72] E. Santiago Mujika, R. Luthi Carter, F. Giorgini, R. N. Kalaria, and E. B. Mukaetova Ladinska. “Tubulin and tubulin posttranslational modifications in Alzheimer’s disease and vascular dementia”. Frontiers in Aging Neuroscience, 13, 730107, 2021.

[73] A. Sferra, F. Nicita, and E. Bertini. “Microtubule dysfunction: A common feature of neurodegenerative diseases”. International Journal of Molecular Sciences, 21(19), 7354, 2020.

[74] X. Hu, H. Zhu, B. Chen, X. He, Y. Shen, X. Zhang, W. Chen, X. Liu, Y. Xu, and X. Xu. “Tubulin alpha 1b is associated with the immune cell infiltration and the response of HCC patients to immunotherapy”. Diagnostics, 12(4), 858, 2022.

[75] J. A. Vizcaino, E. W. Deutsch, R. Wang, A. Csordas, F. Reisinger, D. Rios, J. A. Dianes, Z. Sun, T. Farrah, N. Bandeira, et al. “ProteomeXchange provides globally coordinated proteomics data submission and dissemination”. Nature Biotechnology, 32(3), 223–226, 2014.

[76] R. Hesse, M. L. Hurtado, R. J. Jackson, S. L. Eaton, A. G. Herrmann, M. Colom Cadena, M. Tzioras, D. King, J. Rose, J. Tulloch, et al. “Comparative profiling of the synaptic proteome from Alzheimer’s disease patients with focus on the APOE genotype”. Acta Neuropathologica Communications, 7, 1–18, 2019.

[77] M. C. Chambers, B. Maclean, R. Burke, D. Amodei, D. L. Ruderman, S. Neumann, L. Gatto, B. Fischer, B. Pratt, J. Egertson, et al. “A cross-platform toolkit for mass spectrometry and proteomics”. Nature Biotechnology, 30(10), 918–920, 2012.

[78] D. Klopfenstein, L. Zhang, B. S. Pedersen, F. Ramirez, A. Warwick Vesztrocy, A. Naldi, C. J. Mungall, J. M. Yunes, O. Botvinnik, M. Weigel, et al. “GOATOOLS: A Python library for gene ontology analyses”. Scientific Reports, 8(1), 1–17, 2018.

[79] D. Szklarczyk, A. L. Gable, K. C. Nastou, D. Lyon, R. Kirsch, S. Pyysalo, N. T. Doncheva, M. Legeay, T. Fang, P. Bork, et al. “The STRING database in 2021: Customizable protein-protein networks, and functional characterization of user-uploaded gene/measurement sets”. Nucleic Acids Research, 49(D1), D605–D612, 2021.

[80] P. Shannon, A. Markiel, O. Ozier, N. S. Baliga, J. T. Wang, D. Ramage, N. Amin, B. Schwikowski, and T. Ideker. “Cytoscape: A software environment for integrated models of biomolecular interaction networks”. Genome Research, 13(11), 2498–2504, 2003.

[81] D. Shcherbakov, Y. Teo, H. Boukari, A. Cortes Sanchon, M. Mantovani, I. Osinnii, J. Moore, R. Juskeviciene, M. Brilkova, S. Duscha, et al. “Ribosomal mistranslation leads to silencing of the unfolded protein response and increased mitochondrial biogenesis”. Communications Biology, 2(1), 381, 2019.

[82] P. Kaiser. “Methionine dependence of cancer”. Biomolecules, 10(4), 568, 2020.

[83] J. Ognjenovic and M. Simonovic. “Human aminoacyl-tRNA synthetases in diseases of the nervous system”. RNA Biology, 15(4-5), 623–634, 2018.

