## supplementary note for "Profiling proteome-level amino acid substitutions in Alzheimer’s disease brain tissue"

for

Yu^1,2^*^,^*^∗^

1 Individualized Interdisciplinary Program,

The Hong Kong University of Science and Technology, Hong Kong, China

2 Department of Electronic and Computer Engineering,

The Hong Kong University of Science and Technology, Hong Kong, China

3 Division of Life Science,

The Hong Kong University of Science and Technology, Hong Kong, China

4 Department of Biomedical Engineering, School of Basic Medical Sciences,

Central South University, Changsha, Hunan Province, China

^†^ The authors contributed equally to this paper

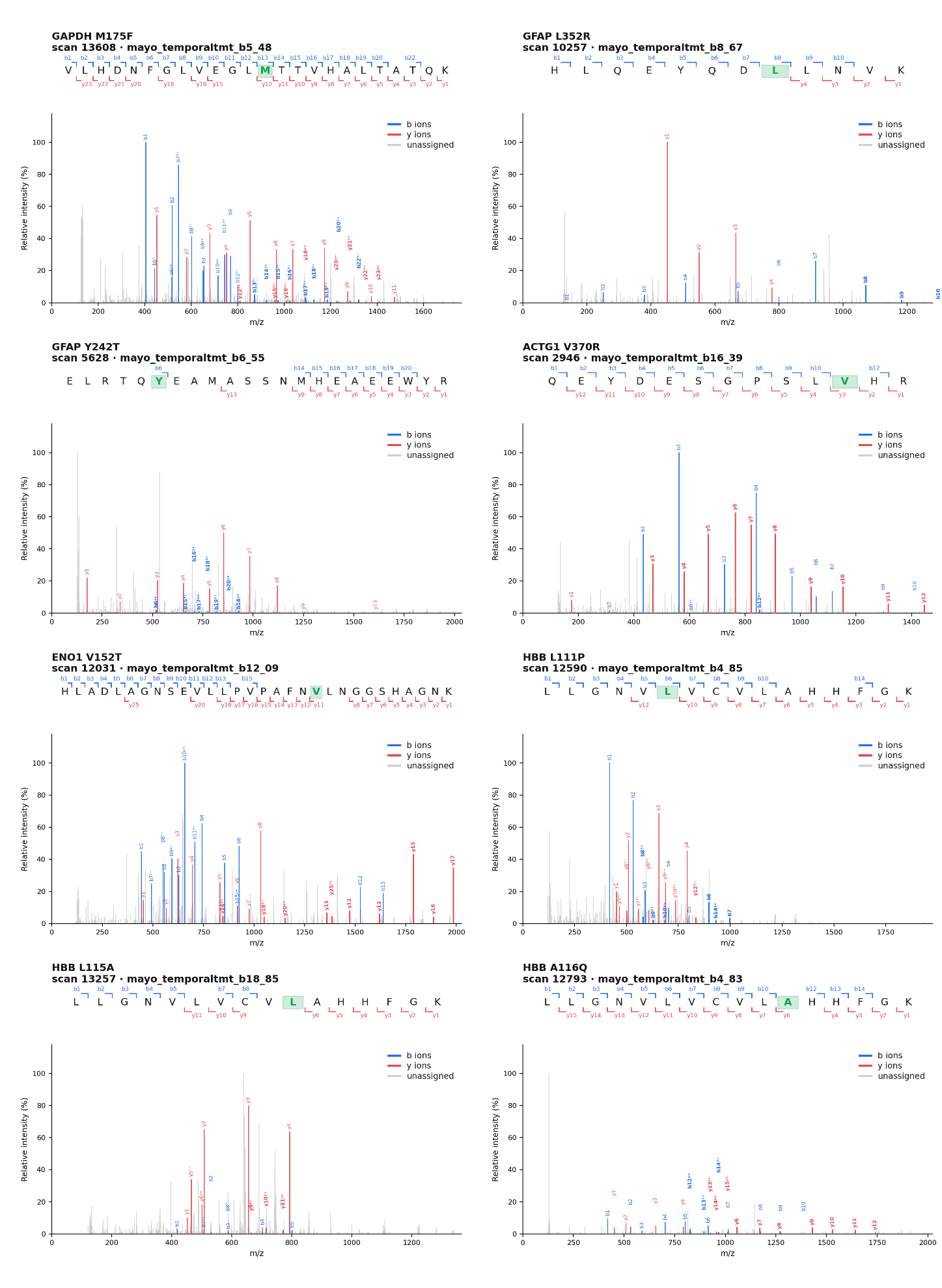

Supplementary Figure S1: Manually annotated MS2 spectra for eight representative AA substitution sites.

For each panel, the upper track shows the peptide sequence with the substituted residue highlighted in green and the matched b (blue) and y (red) cleavage positions marked; the lower track shows the annotated spectrum, with matched b and y ions coloured and unassigned peaks in grey. Ion labels in bold denote site-determining ions, whose sequence span includes the substituted residue and which therefore localize the mass shift to a single position. Panel titles give the source raw file and scan number; the corresponding spectra are available in the Zenodo deposit (record 15010408). Raw file names carry the mayo_temporaltmt prefix used for this deposit; every TMT batch contains specimens from both contributing institutions (Emory and Mayo Clinic) and was acquired at Emory.
